# Looking for a sign forecasting failure in actions: reaching errors triggered by a slowdown of movement and specific brain activity in preceding trials

**DOI:** 10.1101/2022.08.23.505043

**Authors:** Toshiki Kobayashi, Mitsuaki Takemi, Daichi Nozaki

## Abstract

Even experts can sometimes fail while performing fully learned movements. Do such failures suddenly arise, or are there any forecasting signs? It has been reported that the kinematics of the early phase of movements can predict the failure, and brain activity patterns specific to failures are observed just before the movement onset. The presence of abnormal brain activity patterns long before (> 30 s) a failure in a cognitive task leads us to question if signs of a failure in action could exist in trials preceding the failure. Here, we examined this question using a reaching movement adaptation paradigm conventionally used to test motor learning dynamics. Firstly, the presence of a behavioral sign that preceded failures was observed: the peak velocity of the reaching movement significantly decreased in the preceding two trials. Secondly, specific theta and alpha band activity of EEG were observed in the failure trials and the trials preceding the failure. These results suggest that a failure in actions does not occur suddenly, and some signs preceding failures can be observed in the prior trials. Our approach may pave the way to investigate how we prevent failures and improve motor performance.

## Introduction

Humans have a remarkable ability to learn incredibly sophisticated motor skills by repeated practice. However, even after a skill has been learned sufficiently, the occurrence of failure is sometimes inevitable. For example, it has been observed that the failure rate for even top NBA players exceeds 10% on the free throw^1^. Similarly, elite football players fail penalty kicks with a probability of >20%^2^. Failure is often associated with high-pressure situations where successful performance is critically important. Psychological factors including anxiety^3^, diverted attention^4,5^, or excessive focus on movement execution^5,6,7^ could adversely influence the performance and lead to failures. Nieuwenhuys & Oudejans^3^ proposed a theory that in extreme situations that mandate the best performance (e.g., a match awarding prize money or a world-championship final), attentional control focused on task-relevant information is distracted, or automatic execution of well-skilled behavior is disrupted.

The occurrence of failure is, however, not restricted to cases where high pressure is involved. In our daily lives, we also make failures: mistyping words^8,9,10^, driving errors^11^, and stumbling during walking^12,13^. Considering such action failures could sometimes lead to significantly grave outcomes (traffic accidents or bone fracture in the elderly), understanding their mechanisms is very important to avoid them. Previous studies have demonstrated that whether an action is going to fail is already apparent in the early stages of movement, such as in a basketball free throw^14^, piano finger tapping^15^, and ring-throw^16^. These results indicate that a failure may partly arise in the motor planning stage.

In accordance with these behavioral studies, several previous studies have observed brain activity patterns specific to failure trials (the trials where failure was achieved) before the movement onset. For example, Ruiz et al.^15^ found that around 70 ms prior to an erroneous key-press during a piano performance, the event-related potential at the frontocentral brain regions became weak. Brain activity preceding an error has also been reported during Go/NoGo tasks. Bediou et al.^17^ demonstrated that incorrect responses to NoGo cues were associated with larger amplitudes of event-related potential over the medial frontal region 100 ms before the responses. Babiloni et al.^18^ also showed that the weaker the attenuation of the alpha band power, the larger the error of the unsuccessful golf putts, over the frontal midline and the upper limb region of the right primary sensorimotor area, 1 s before movement onset.

These previous studies focused on brain activity just prior to the onset of movement, but several other studies have demonstrated that brain activity predicting the failure arises much earlier^19,20,21,22,23^. For example, a functional MRI study, applying an independent component analysis with a cognitive task, revealed that a specific set of brain regions had linear trends starting from >30 s before erroneous responses occurred^22^. Hence, we arrive at the following questions: can changes in brain activity, long before the failure trials, also be observed in sensorimotor tasks? Are the changes accompanied by any observable change in the movement patterns? In order to answer these questions, this study adopted a reaching adaptation task to a novel force field, which has been conventionally used to investigate motor learning behavior^24,25,26,27^. As demonstrated in the following sections, even after participants fully adapted to the force field, greater movement errors were still occasionally exhibited by them. Thus, this task could allow us to explore the possible signature of movement patterns and brain activity preceding the occurrence of movement errors.

## Results

### A behavioral sign of failure in action

In Experiment 1, 15 young, healthy participants (22–24 years old, right-handed) performed 400 reaching movements towards a frontal target (movement distance 10 cm) with their right arm in the presence of a clockwise (CW) velocity-dependent force field (Fig. 1a). The reaching error evaluated by the lateral deviation of the handle trajectory at the peak velocity gradually decreased with the training trials. After 100 training trials, the reaching error seemed to converge to a steady-state level. However, even after the completion of the training, the reaching errors varied and exhibited sudden greater deviations in certain trials (Fig. 1b). The trials with the top 5% error values (i.e., worst 15 trials) were defined as the failure trials (FTs).

**Figure 1.**
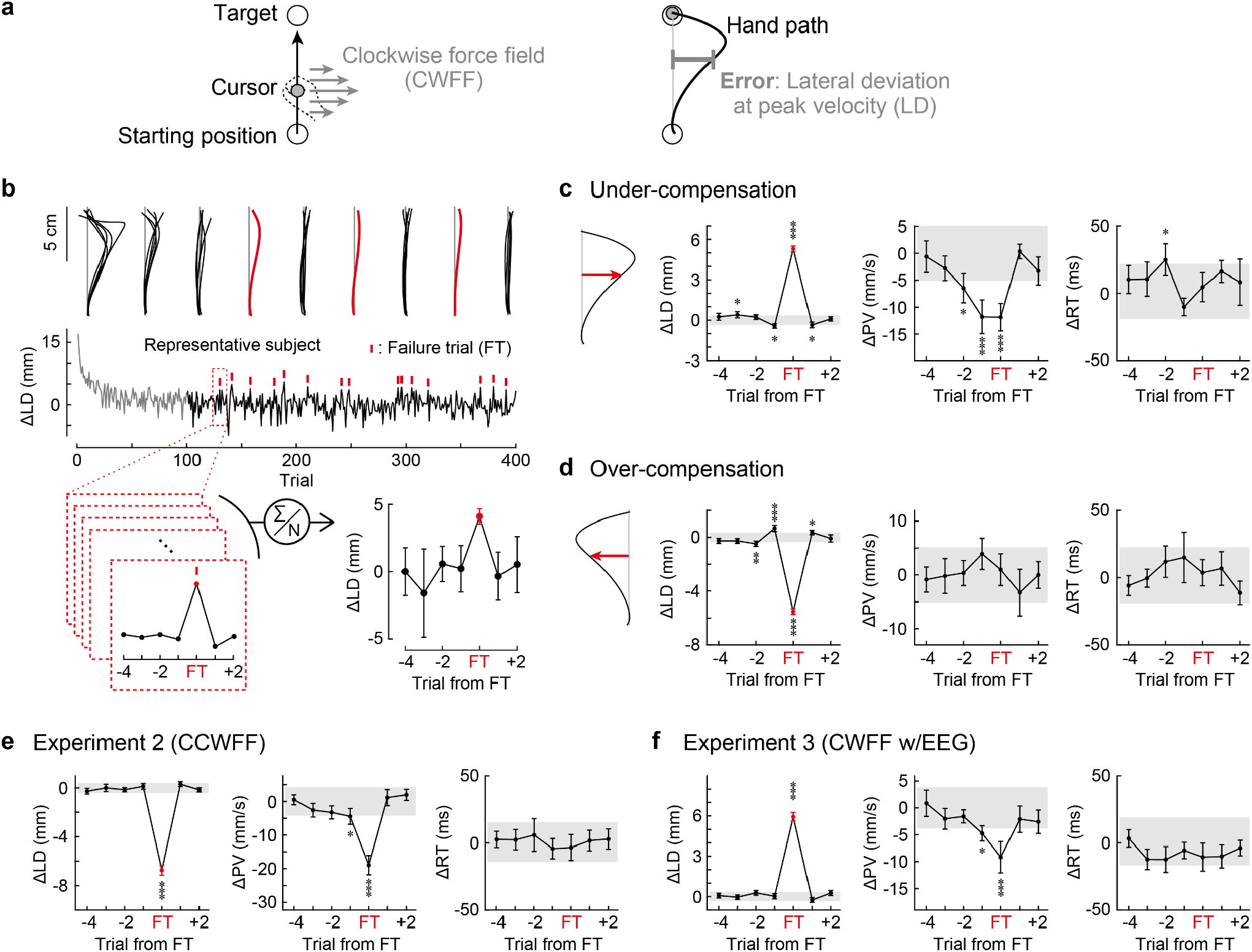
Searching for a sign preceding the failure. (**a**) A clockwise velocity-dependent force field (CWFF) was imposed on a handle during reaching movement. Reaching error was defined as the lateral deviation (LD) between an ideal trajectory (gray straight line) and the cursor path (black line) at peak velocity. (**b**) A procedure to search for a sign preceding the failure from the data of a representative subject. *Top:* Cursor trajectories of the subject. Even after completion of the training, the reaching errors exhibited sudden deviations for certain trials (red line). *Middle:* The trial-dependent change in ΔLD during the task. The trials in which the top 5% values of error (i.e., worst 15 trials) were observed were defined as the failure trials (FTs). *Bottom:* The data around the FT (4 trials before to 2 trials after the FT) were selected and averaged to observe the change in the behavioral parameters. (**c**) Averaged data of the error in the force field direction (i.e., under-compensation error), the peak velocity (PV), and the reaction time (RT) across subjects (\**P* < 0.05, \*\**P* < 0.01, \*\*\**P* < 0.001, by bootstrap resampling test). Error bars denote the SEM. (**d**) The over-compensating error in the direction opposite to the force field direction did not show any sign of forecasting the FTs. (**e, f**) The behavioral results of Experiment 2 and 3, respectively.

Next, the data around the FT (4 trials before to 2 trials after the FT) were considered, and the data was averaged to observe changes in the behavioral parameters around the FT (Fig. 1b). There was no specific trial-dependent trend in reaching errors nor in reaction time in trials preceding the FT (Fig. 1c), although significant differences were observed in a few trials (e.g., 3 trials before the FT in lateral deviation). However, the peak movement velocity was found to have already started declining from two trials preceding the FTs (*P* < 0.05, compared to resampled trials) (Fig. 1c).

Notably, the slowdown of the movement velocity was observed only in the case of under-compensation errors. Any signature in the peak velocity profile in the trials preceding the over-compensating errors were not found (Fig. 1d). Thus, the slowing down of the peak velocity was specific to the trials preceding the under-compensating errors. Considering that the inter-trial interval (ITI) of this experiment was 4.5−5.0 s, the two trials corresponded to approximately 10 s before the FTs. Hereafter, the trials with under-compensating errors are consistently referred to as the FTs.

### Replication of the behavioral results

The gradual decline of peak velocity before the FTs was obtained by exploring the data in Experiment 1. To examine whether this result can be replicated, 15 new participants were recruited (Experiment 2). In this experiment, the direction of the force field was changed to the counter-clockwise (CCW) direction. The results were substantially similar to Experiment 1 (Fig. 1e): The FTs suddenly occurred, and the reduction of the peak velocity was already observed in the trial before the FT (*P* < 0.05, compared to resampled trials). Lastly, Experiment 3 was performed to investigate if such a sign of failure that was observed even before the FTs could be detected in brain activity by measuring EEG (N = 15). The behavioral results again replicated the observations of the previous experiments: The sudden occurrence of FTs and the decrease in the peak velocity in the trials before the FTs were observed (Fig. 1f, *P* < 0.05, compared to resampled trials). However, in Experiment 2 and 3, a significant decrease in the peak velocity was observed only for the trial that was the immediate predecessor of the FT, which was possibly due to the experiment setting in which the ITI was lengthened (Experiment 1: 4.5−5.0 s, Experiment 2: 4.5−6.5 s, Experiment 3: 6.5−8.5 s; In Experiment 3, the ITI needed to be considerably lengthened not to contaminate EEG activity by the prior trials). Despite these slight differences, the consistent empirical results strongly demonstrated that the behavioral sign of the FTs could be detected at 1−2 trials before the FT.

### Influence of the definition of the failure trials

In these behavioral results, a threshold was adopted to determine the FTs as the worst 5% trials. Since this threshold value was arbitrary, an examination was conducted to determine the influence of different thresholds on the results. The results were largely unaffected when this percentage threshold in the definition of FTs was varied from 1% to 17% (Fig. 2, calculated using all behavioral data of Experiments 1–3). For example, when the worst 10% trials were defined as the FTs, the peak velocity was significantly slower in the 1^st^ preceding trial (−5.3 ± 1.2 mm/s relative to average velocity, *P* < 0.001, compared to resampled trials). The trend of the slowing down of the velocity attenuated gradually as the percentage value in the definition of the FTs increased. Note that when the worst 15% trials were defined as the FTs, the slowing down of the velocity in the 1^st^ preceding trial was significant (−4.1 ± 1.0 mm/s relative to average velocity, *P* < 0.01, compared to resampled trials) but not in the 2^nd^ preceding trial (−1.0 ± 0.7 mm/s relative to average velocity, *P* = 0.445, compared to resampled trials).

**Figure 2.**
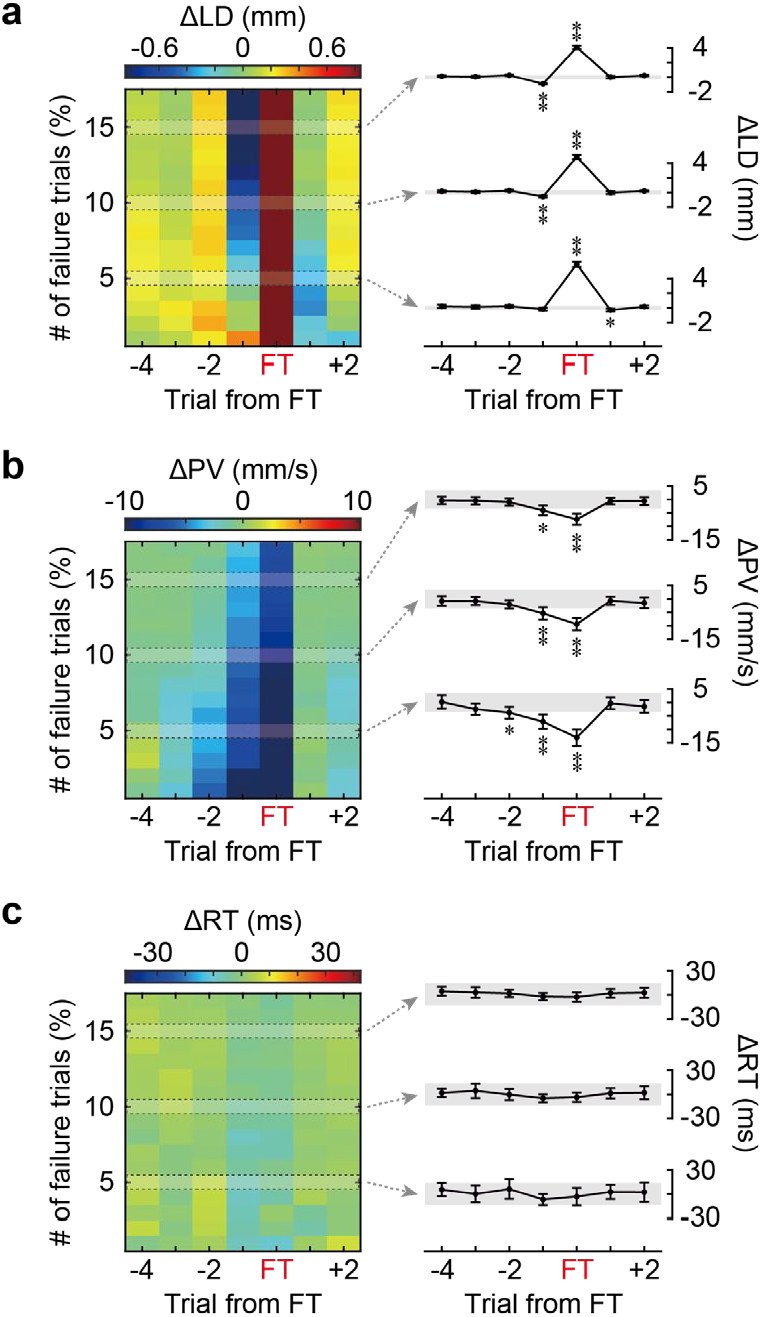
Influence of the definition of FTs. (**a**) The left colormap illustrates how the definition of FTs (i.e., the percentage of trials considered as FTs) affects the ΔLD in the trials before and after the FTs. The data from Experiments 1–3 were analyzed. The results for 5%, 10%, and 15% were plotted on the right with the same format as Fig. 1c. (**b, c**) PV and RT. The slowing down of the movement preceding the FTs was substantially unaffected when the definition of FT was changed from the worst 1 % to 17 %. \**P* < 0.01, \*\**P* < 0.001, with the bootstrap resampling test.

### Random occurrence of the failure trials

Did the FTs occur in a specific phase (early or late) of the experiments? Figure 3a indicates how the FTs were distributed in the last 300 (Experiment 1) or 400 adaptation trials (Experiment 2 and 3). The occurrence ratio of the FTs was calculated for each participant in 4 adaptation phases (Fig. 3b). If the FTs occurred uniformly, then the ratio should be around 5% for all 4 phases. A repeated-measures ANOVA indicated that there was no significant difference in the occurrence ratio among the 4 phases (*F*(3, 142) = 1.1587, *P* = 0.3281), indicating there was no specific phase where the FTs occurred more frequently (e.g., late phase due to fatigue or the lack of concentration). This result was moderately supported by a complementary Bayesian repeated-measures ANOVA analysis (*B*_*10*_ = 0.1799).

**Figure 3.**
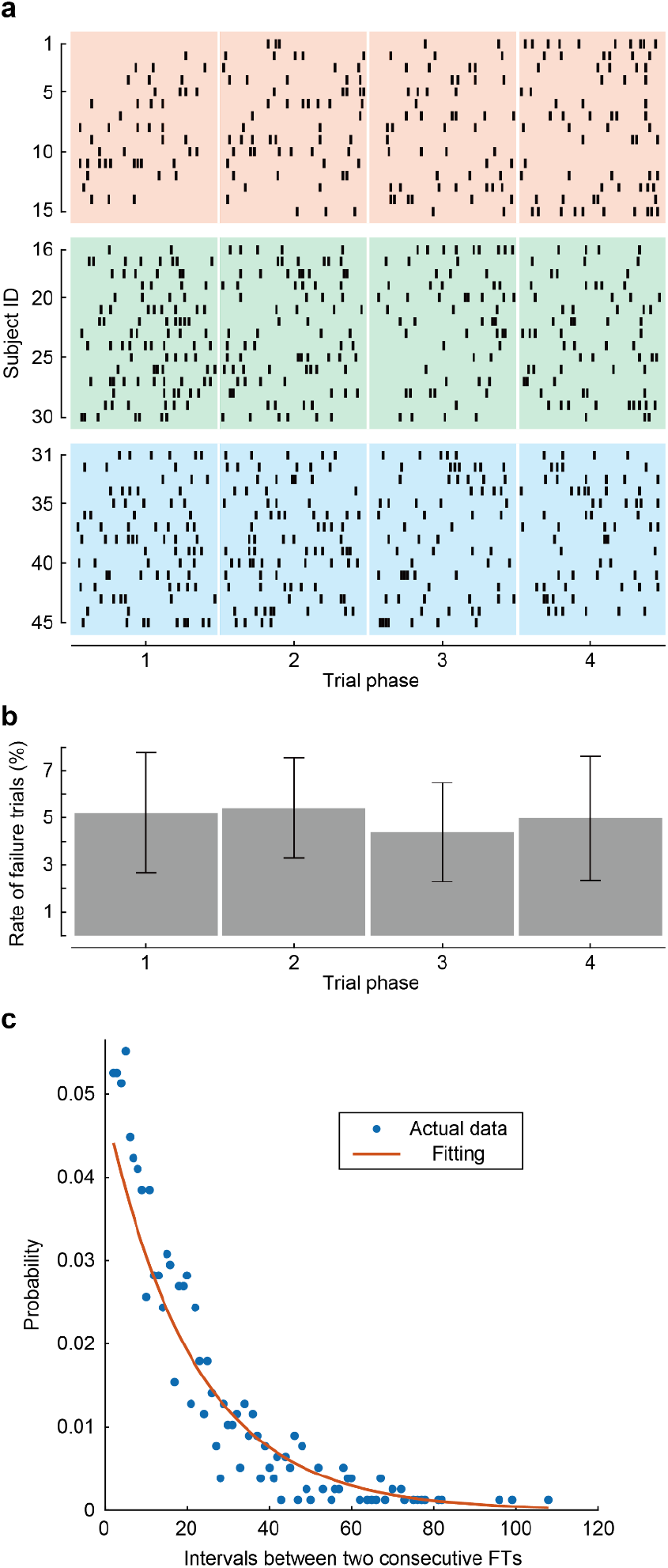
The randomness of the occurrence of the FTs. (**a**) Distribution of the FTs in the last 300 (Experiment 1) or 400 adaptation trials (Experiment 2 & 3). Each bar indicates the timing (trial) at which an FT occurred. Subject ID 1–15, 16–30, and 31–45 indicate subjects in Experiment 1, 2, and 3, respectively. (**b**) The occurrence ratio of the FTs was calculated for each participant in 4 adaptation phases. Error bars denote the SD. A repeated-measures ANOVA indicated no significant difference in the occurrence ratio among the 4 phases. (**c**) The distribution of intervals between two consecutive FTs (blue dots) was fitted to an exponential distribution *f*(*T*) = *λ*exp(−*λ* (*T* − 1)) (red line), where *T* is the interval between two consecutive FTs. The obtained value was close to the theoretical value of 0.05 (the definition of FT).

Furthermore, to examine how the FTs occurred randomly, the distribution of intervals between two FTs was analyzed. If the FTs occurred randomly, the distribution of intervals between two FTs (*T*) was expected to follow an exponential distribution^28^. The distribution was fit best by an exponential function *f*(*T*) = *λ*exp(−*λ* (*T* − 1)) with *λ* = 0.047 (Fig. 3c). The value of 0.047 was close to the theoretical value of 0.05 (the definition of FT). Note that a part of the equation of the exponential distribution was modified because the second and subsequent trials of consecutive failure trials were not counted as failure trials (i.e., T = 1 does not exist).

### Brain activity around failure trials

As we described above, in Experiment 3, we observed the behavioral sign in the preceding trial of the FT. To investigate if a specific electroencephalographic activity was associated with the failure, a time-frequency analysis was conducted at eight time-locked epochs (Fig. 4a): 0.5 s before and after the target cue, the movement cue, the movement onset, and the movement end. The power of each time epoch was normalized to the power of the baseline period (1 s before the target cue), expressed as the event-related (de)synchronization (ERD/ERS)^29^. Topographic representations of ERD/ERS were analyzed using a cluster-based permutation testing to circumvent the multiple comparisons problem between channels^30^. Interestingly, as shown below, the behavioral signature was likely to be associated with brain activity during the movement preparation period.

**Figure 4.**
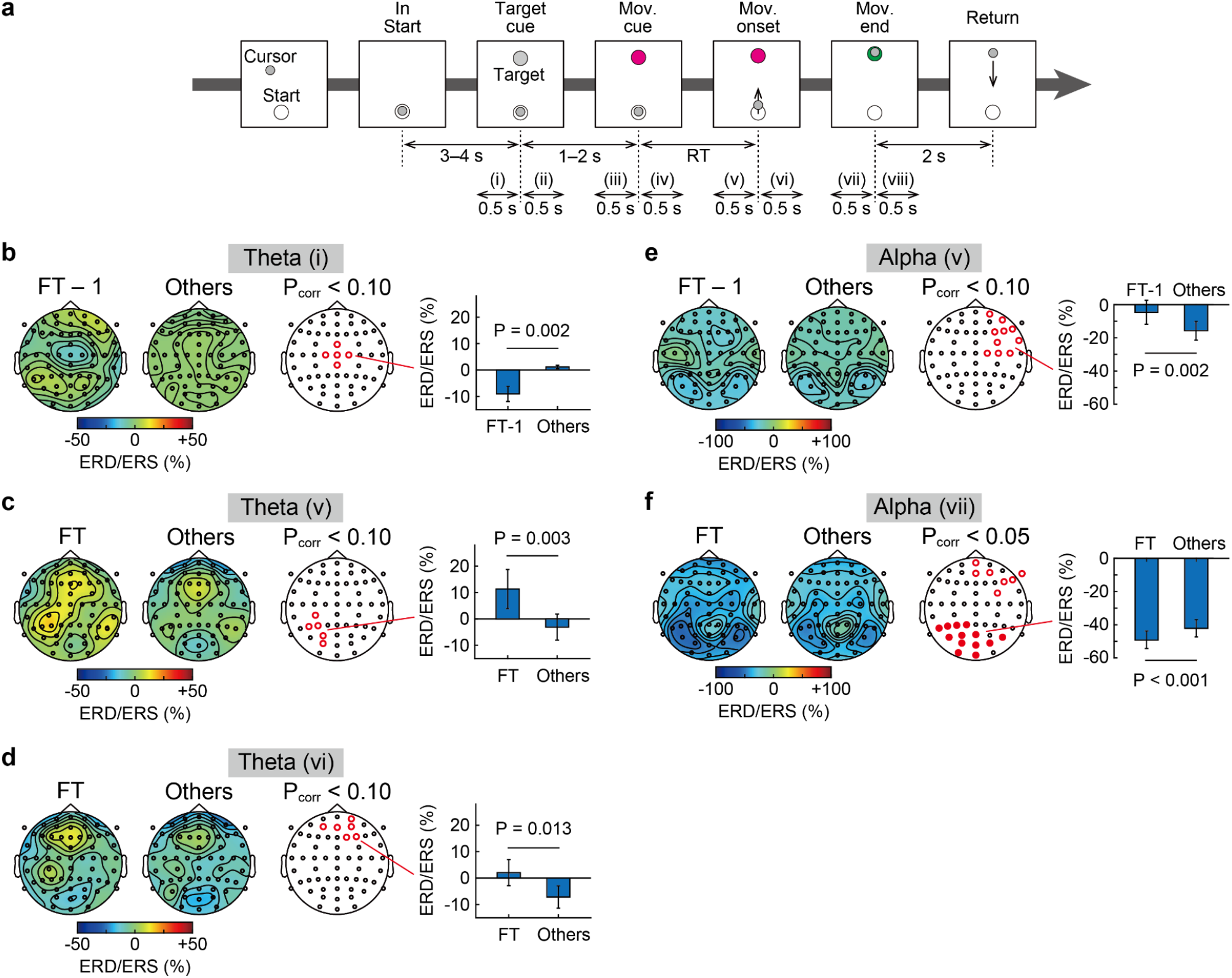
Electroencephalographic activities associated with the failure. (**a**) A trial was divided into eight time-epochs. (**b**) Left and middle topographic maps show the theta ERD/ERS at the time-epoch (i) in the trials preceding the FTs and in the other trials, respectively. The right topographic map shows statistical results by cluster-based permutation testing. Red rings denote *P* < 0.10. Averaged ERD/ERS across the channels in the detected cluster was compared using a paired *t*-test (the right bar graph). (**c**–**d**) The theta ERD/ERS at the time-epoch (v) and (vi) in the FTs. (**e**) The alpha ERD/ERS at the time-epoch (v) in the trials preceding the FTs and (**f**) at the time-epoch (vii) in the FTs. Red filled circles denote *P* < 0.05 with cluster-based permutation testing.

The first area of focus was the theta band (4–7 Hz) activity that reflects attentional control and is a factor related to the failure^31,32,33^. The results demonstrated that the theta ERD/ERS significantly shifted in the period prior to the failures, but its trend was not consistent (Fig. S1). In the trial preceding the FT, the power in the midline sensorimotor area was significantly attenuated for 0.5 s before the target cue (*P* < 0.10 by the cluster-based permutation testing, Fig. 4b). Meanwhile, in the FT, a larger theta power was observed over the contralateral frontoparietal regions 0.5 s before the movement cue and the movement onset (*P* < 0.10 by the cluster-based permutation testing, Fig. 4c, d).

The alpha band (8–13 Hz) ERD/ERS was also analyzed, which showed less power decrease just before the movement in the failure trials^18^. The alpha ERD/ERS showed the opposite trend to the theta ERD/ERS. In the trial preceding the FT, the ERD in the ipsilateral frontal area was weaker for 0.5 s before the movement onset (*P* < 0.10 by the cluster-based permutation testing, Fig. 3e). In the FTs, the ERD in the occipital-parietal area was greater for 0.5 s before the movement end (*P* < 0.05 by the cluster-based permutation testing, Fig. 4f).

## Discussion

In this study, a slowing down of the movement in at least one trial preceding the failure trials (FTs) was discovered by three independent experiments using the velocity-dependent force field. EEG measurements during the motor task also indicated changes in the brain activity before the movement onset in one trial immediately preceding the FTs. These results suggest that the FTs do not suddenly occur without any sign. Instead, they are preceded by the signs of the behavior and brain activity.

### Definition of the failure trials

First, the validity of our approach needs to be discussed. In the present study, the FTs were defined as the trials that constituted the worst 5% movement errors at the peak velocity. However, even if such a larger error was produced, the participants were able to correct the movement through online feedback. Thus, the larger error does not imply that the trials ended unsuccessfully. Additionally, the participants were scarcely aware of the error occurrence. From these viewpoints, the FTs defined in this study did not directly correspond to ordinary failure (e.g., missing the target). This is the reason why slowing down of the movement in the trials following the FT (i.e., post-error slowing)^8,15,34,35,36^ was not observed in this study (Fig. 2b). However, this is not a significant problem because the main focus here was on the trials preceding the failure trial. Furthermore, considering that the awareness of error leads to explicit movement correction^37^, which might often produce greater movement variability^38^, our approach is believed to be useful for precisely capturing the unintentional process of producing the movement error.

### Kinematical signs of failure trials

The trial-by-trial change in the movement error can be described by the state-space model in which the motor command is updated by the movement error^24^. According to this idea, a greater (i.e., under-compensated) error should be preceded by a smaller error or by an error in the opposite direction (i.e., over-compensation). This tendency existed when the FTs were defined as the worst 15% trials (Fig. 2a). However, as the criteria for an FT became more severe (e.g., 5%), the tendency became more obscured (Fig. 1; Fig. 2a) and the FTs appear to occur suddenly. In other words, the movement error in FT-1 was not small enough to produce a greater error in FT, and the movement error was unlikely to have sufficient information to predict the occurrence of an FT.

Instead, our results indicate that the movement velocity had information to predict the future FT. Notably, slow movements were observed not only in the FT-1 but also in the FTs (Fig. 1c; Fig. 2b). One possible factor slowing down the movement in these trials is fatigue and/or lack of concentration induced by the prolonged adaptation trials. In that case, the number of FTs should increase as the experiment progresses. However, no significant difference was observed in the occurrence rate of FT in the 4 different phases (Fig. 3b), indicating that fatigue and/or lack of concentration by the prolonged adaptation trials were unlikely to cause the FT.

We speculate that the factor contributing to the slowdown of the movements preceding the FTs is related to the temporal reduction of movement vigor and implicit motivation. Previous studies have shown that reduced movement vigor and implicit motivation slowed movement^39,40,41^. Additionally, it has been shown that as a reward decreases, vigor decreases and motor performance worsens^42,43^. The decrease in movement velocity in our results can be attributed to the reduction of vigor and motivation, resulting in slower movement and large motor errors. This idea is also consistent with the result that the over-compensated error was not preceded by slow movement (Fig. 1e). Zhu et al.^32^ reported that lower vigor and motivation are involved in exerting more attenuated hand grip force, which is also consistent with the factor of the under-compensating error in our study (i.e., insufficient hand force to counteract the force field).

### Signs of failure trials in EEG activity

Cavanagh et al.^31^ reported that attentional lapses were observed in the trial preceding failure trials with weaker theta power in the medial prefrontal cortex. Similarly, our results also showed weaker theta power in the midline sensorimotor area in the trial preceding FTs. Additionally, a larger theta power in the parietal area during movement planning in the FTs in our results may reflect lower vigor and motivation^32^. Our results also showed larger theta power in the frontal area following the movement onset, which may reflect the increased attention induced by implicit error detection.

The results of this study demonstrated less decrease in the alpha power relative to the baseline in the trial preceding the FTs. In line with a previous study showing that an increase in alpha power precedes errors in attentional tasks^23^, these observed changes in the alpha ERD/ERS may reflect attentional lapses^33,44,45^. Additionally, decrease in alpha power in the FTs during movement execution may reflect alpha suppression, known to be observed at the timing of implicit error detection^33,44,45^. Taken together, both theta and alpha oscillations before errors are considered to be associated with reduced vigor and motivation through attentional lapses^46^.

In summary, the present study revealed the presence of signs in the trials before the failure trial. The slowing down of the movement was observed by exploring the kinematics before the failure trials, and the results were then replicated. In addition, such behavioral signs were also accompanied by specific brain activities. We assume that this sign is related to the reduction of vigor and motivation, but it is possible that the other motor tasks could have different behavioral or neuronal signs. However, we believe that our approach helps investigate the universal mechanism of producing the failure trials and detect the failure trials beforehand to prevent failures.

## Acknowledgement

We thank members of the Nozaki laboratory for their helpful comments and suggestions, Naoki Hashimoto for performing preliminary experiments, Asako Munakata for coordinating experiments. This study was supported by a grant from the Japan Society for the Promotion of Science Research Fellowships for Young Scientists to T.K. (20J13734), JST PRESTO to M.T. (JPMJPR18J6) and a KAKENHI to D.N. (17H00874, 21H04860).

## Author contributions

Conceptualization, Methodology, Investigation, Formal Analysis, Writing, T.K., M.T., D.N. Supervision, D.N.

## Materials and Methods

### Participants

A total of 45 healthy adults participated in the study (Experiment 1: 15 males, aged 22−24 years, all subjects were right-handed according to self-reports. Experiment 2: 8 females and 7 males, aged 23.3 ± 6.6, laterality score (LS): 87.3 ± 12.4; Experiment 3: 4 females and 11 males, aged 25.1 ± 3.2, LS: 65.3 ± 34.2; LSs were derived from the Edinburgh Handedness Inventory^47^). All subjects gave written informed consent to an experimental protocol approved by the ethical committee of The University of Tokyo (#18-203).

### Experimental apparatus

The participants performed a task with a kinesiological instrument for normal and altered reaching movements (KINARM End-Point Lab, BKIN Technologies, Kingston, Ontario, Canada)^48,49^. The robotic manipulandum allowed planar hand movements and could apply independent mechanical loads to the hand. The subjects were seated in front of an LCD monitor projecting visual targets and hand-aligned feedback, referred to as the ‘cursor’ (10 mm diameter circle), and a direct vision of their limb was occluded. The position and velocity of the hand were analog/digital converted at 1.129 kHz and then recorded at 1 kHz for subsequent offline analysis.

### Experiment 1: searching for a behavioral sign

This study adopted a motor adaptation task using a velocity-dependent force field. There are several reasons for choosing this task over alternative tasks such as visual rotation^50^ or divergent force field^51^. First, the task produced a moderate size error at a moderate frequency. In contrast, the error size after the participants adapted to the visual rotation task was too small, and the failure frequency was too often for the divergent force field. Second, contribution of the explicit component was to be minimized because the explicit component often causes greater movement variability and error^38^. Compared to the visual rotation or divergent force field tasks, the contribution of the velocity-dependent force field task was small (but not negligible)^52^.

Fifteen participants grasped the robotic manipulandum handle with their right hand and moved it from a home position (14 mm diameter circle) toward a forward target (14 mm diameter circle) that was 10 cm away from the home position (Fig. 1a). After 10 null trials for practice, the participants repeated arm-reaching movements in the presence of a clockwise velocity-dependent force field (400 trials). The force, *f* = (*f*_*x*_, *f*_*y*_) (N), imposed on the handle was always set to be perpendicular to the velocity of the handle, *v* = (*v*_*x*_, *v*_*y*_) (m/s), as *f* = *Bv*, where *B* = (0 15; -15 0) [N/(m/s)]. In each trial, after the subjects kept the cursor at the home position for 2.5−3 s, the target appeared on the monitor, indicating that they should immediately start moving the cursor to the target (i.e., movement cue). The subjects were instructed to maintain the peak velocities as constant as possible across the trials. When the cursor reached the target, a feedback message, ‘fast’ or ‘slow’, was presented on the monitor if the movement speed was faster or slower than the range of 250−450 mm/s, respectively. After maintaining the position of the cursor at the target for 2 s, the handle automatically returned to the home position, and the next trial began.

### Experiment 2: validation of the results of Experiment 1

Based on the results of Experiment 1, we found that the FTs were preceded by the trial(s) where the peak hand velocity slowed down. Experiment 2 was performed with 15 newly recruited participants to investigate if these exploratorily obtained results could be replicated. In this experiment, the direction of the force field was opposite to that of Experiment 1 (i.e., counter-clockwise direction: *B* = (0 -15; 15 0) [N/(m/s)]). In addition, extra waiting time was added as the preparation for reaching between staying at the home position and the movement cue. After subjects kept the cursor at the home position, the gray target appeared on the monitor, indicating that they should check the target position and prepare to move the cursor. After maintaining the position of the cursor at the home position for 1−2 s, the color of the target changed to pink, which signaled the participants to move the cursor to the target (movement cue). Thus, the net inter-trial interval was 0–1.5 s longer as compared to that of Experiment 1. The total number of trials was increased to 610 trials to collect a sufficient amount of behavioral data. A short break (2–3 min) was taken every 200 force field trials.

### Experiment 3: searching for an electroencephalographic sign

To examine if the behavioral signs observed in Experiments 1 and 2 were associated with any specific brain activity patterns, we conducted Experiment 3 (N = 15) which took electroencephalograms (EEGs) while participants performed reaching movements in the presence of force field perturbation. The starting position, the target position, the applied force field, and the criteria for the hand velocity feedback were the same as Experiment 1.

The participants performed 10 null trials, the data of which were omitted from further analyses, and then 600 force field trials. A short break was taken every 200 force field trials. We also modified the temporal sequence of a single trial in the same way as Experiment 2. Furthermore, we lengthened the overall inter-trial interval in order to properly investigate preparatory EEG activity and to avoid the possible influence of the post-movement activity of the preceding trial on the baseline EEG. After participants kept the cursor at the home position for 3−4 s, the gray target appeared on the monitor for 1−2 s (target cue). Thereafter, the color of the target changed to pink (movement cue). Participants were instructed to maintain the handle peak velocities as constant as possible across the trials, and the visual feedback of handle velocity was provided (if needed) when the cursor reached the target. After keeping the cursor at the target for 2 s, the handle automatically returned to the home position and the next trial began.

While the participants performed the reaching movement task, EEG signals were continuously recorded using 60 scalp electrodes mounted on a cap according to the 10-20 layout (EASYCAP, Herrsching, Germany) at a sampling rate of 500 Hz using 16-bit biosignal amplifier (BrainAmp DC, BrainProducts, Gilching, Germany). The electrical reference was located at the right earlobe, and the ground electrode was located at the forehead. Electrodes were thoroughly prepared using Nuprep Skin Prep Gel (Weaver & Co., Aurora, Colorado, USA) and Abralyt HiCl EEG electrode gel (Easycap). The impedance of all electrodes was kept below 5 kΩ during all recordings.

### Behavioral data analysis

Behavioral data analysis was performed using MATLAB R2021a (Mathworks, Natick, Massachusetts). Any lateral deviation from the straight path between the target and the home position at the peak hand velocity was considered as a reaching error, i.e., errors whose direction was the same as the force field were considered as the reaching errors because this insufficient compensation implies failures in the adaptation. The trials contributing to the top 5% of reaching errors during the well-practiced phase, i.e., 101−400th force field trials for Experiment 1 and 201−600th force field trials for Experiment 2 and 3, were defined as failure trials. To look for a sign of failures in action, lateral deviation (i.e., the reaching error), peak velocity, and reaction time in the trials preceding and following the failure trials were analyzed. Outliers above four standard deviations were winsorized. To eliminate inter-individual variability, the mean value of the well-practiced phase was subtracted from the result of each subject (e.g., ΔLD). Note that the data of the first 4 trials following the short break (Experiment 2 and 3), where the lateral deviation of the hand trajectory temporarily increased, was omitted. Additionally, the second and subsequent trials of consecutive failure trials were not counted as failure trials. Next, the data around the failure trials (4 trials before to 2 trials after the failure trials) were examined to observe the change in the behavioral data. No subject was excluded from the analysis.

The behavioral data was statistically evaluated by bootstrap resampling^53^. Random trials (5% of overall trials) from each subject’s well-practiced phase data were extracted and then averaged. This procedure was repeated 10,000 times, and then the obtained distribution was used to determine the significance level (i.e., a confidence interval of 95% indicates a range between the 250th largest value and the 9750th largest value). A repeated-measures ANOVA and Bayesian analysis were performed to test the rate of occurrence of the failure trials using the software package JASP (https://jasp-stats.org).

### EEG data analysis

Time-frequency analysis for EEG data was performed using MATLAB R2021a and Fieldtrip toolbox^54^. The raw EEG signals were first filtered with a 1−100 Hz bandpass filter and a 50 Hz notch filter and then re-referenced to the average signal across all electrodes. Independent component analysis was then applied to remove components reflecting artifacts (e.g., eye blink, eye movement)^55^. Data were segmented into epochs from -5 to 3 s relative to the movement cue. Artifact-free EEG epochs were decomposed into time-frequency representations in the 3–40 Hz range (frequency step: 1 Hz, time step: 25 ms). A 7-cycle Morlet wavelet was used for the continuous wavelet transformation. A trial was separated into eight periods based on four events: target cue, movement cue, movement onset, and movement end. The periods were defined as -0.5 to 0 s prior to and 0 to 0.5 s subsequent to these four events. At each single frequency and time point, the power exceeding the mean ± 2SD across trials was treated as an outlier, and they were linearly interpolated using the function *interp1* in MATLAB R2021a. ERD/ERS was then calculated for each individual using the following equation

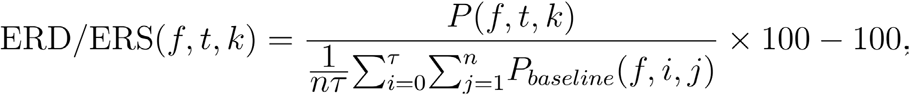

where *P, f, t*, and *k* represent power, frequency, time point in the epoch, and trial, respectively. *τ* and *n* represent the number of epochs in the baseline period (−1 to 0 s prior to the target cue) and the number of trial, respectively (*τ* = 20, *n* = 400). To extract the reactive alpha band, an individual alpha frequency (IAF)^18^ was identified as the frequency showing the strongest ERD (i.e., largest power decrease) within 8–12 Hz at electrode C3 for 500 ms posterior to the movement onset. The alpha band in our study was defined as IAF ± 1 Hz.

A cluster-based permutation testing was conducted for the topographic results to deal with multiple comparison problems between channels^30^. A paired *t* test comparing averages of the FTs (or the trials preceding the FTs) versus averages of the other trials (trials which were neither FTs nor the trials preceding the FTs) was conducted for the logarithm of the rescaled power at each electrode. *t* values exceeding *a priori* threshold of *P* < 0.05 were clustered based on neighboring electrodes. Cluster-level statistics were calculated by taking the sum of *t* values within every cluster. Next, to obtain a null distribution, a permutation of this process was run (i.e., randomizing dataset across trials, rerunning the statistical test 5000 times, and storing the maximum value of summed cluster *t* values). The value at 5% of this distribution (i.e., 250th largest value) was taken as the cluster-corrected threshold. Lastly, the cluster *t* values in the original dataset were evaluated to determine whether they exceeded the corrected threshold.

## Supplementary Figures

**Figure S1.**
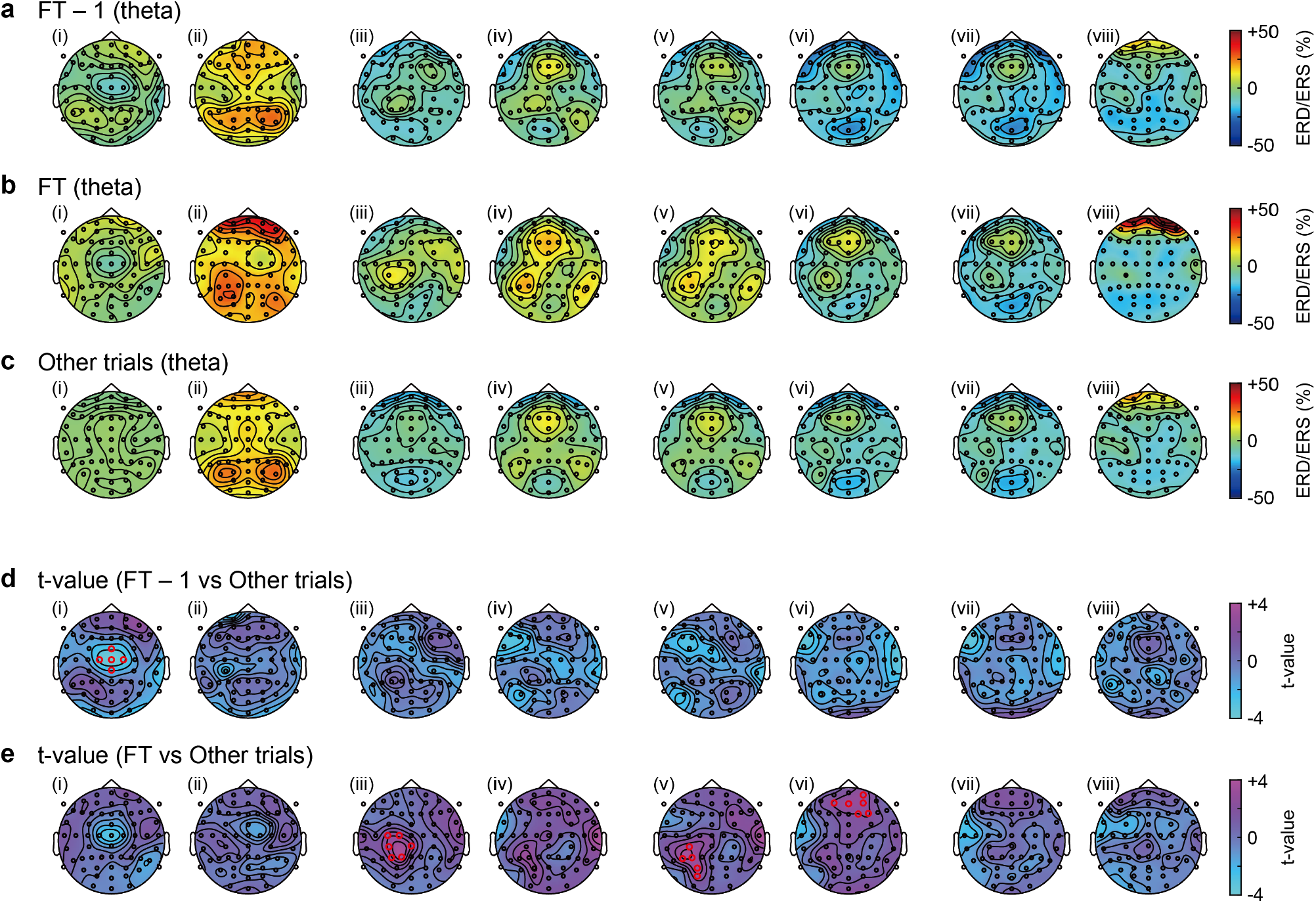
Topographic maps of the theta ERD/ERS. (**a**) ERD/ERS in the trials preceding the FTs, (**b**) in the FTs, and (**c**) in the other trials averaged across subjects. i–viii indicate the time-epoch number in Fig. 3a. ERD/ERS in the FTs averaged across subjects. (**d**) In comparison with the ERD/ERS of the other trials, *t*-values obtained by paired *t*-tests for the trials preceding the FTs, and (**e**) in the FTs are shown. Red rings denote *P* < 0.10 with cluster-based permutation testing.

**Figure S2.**
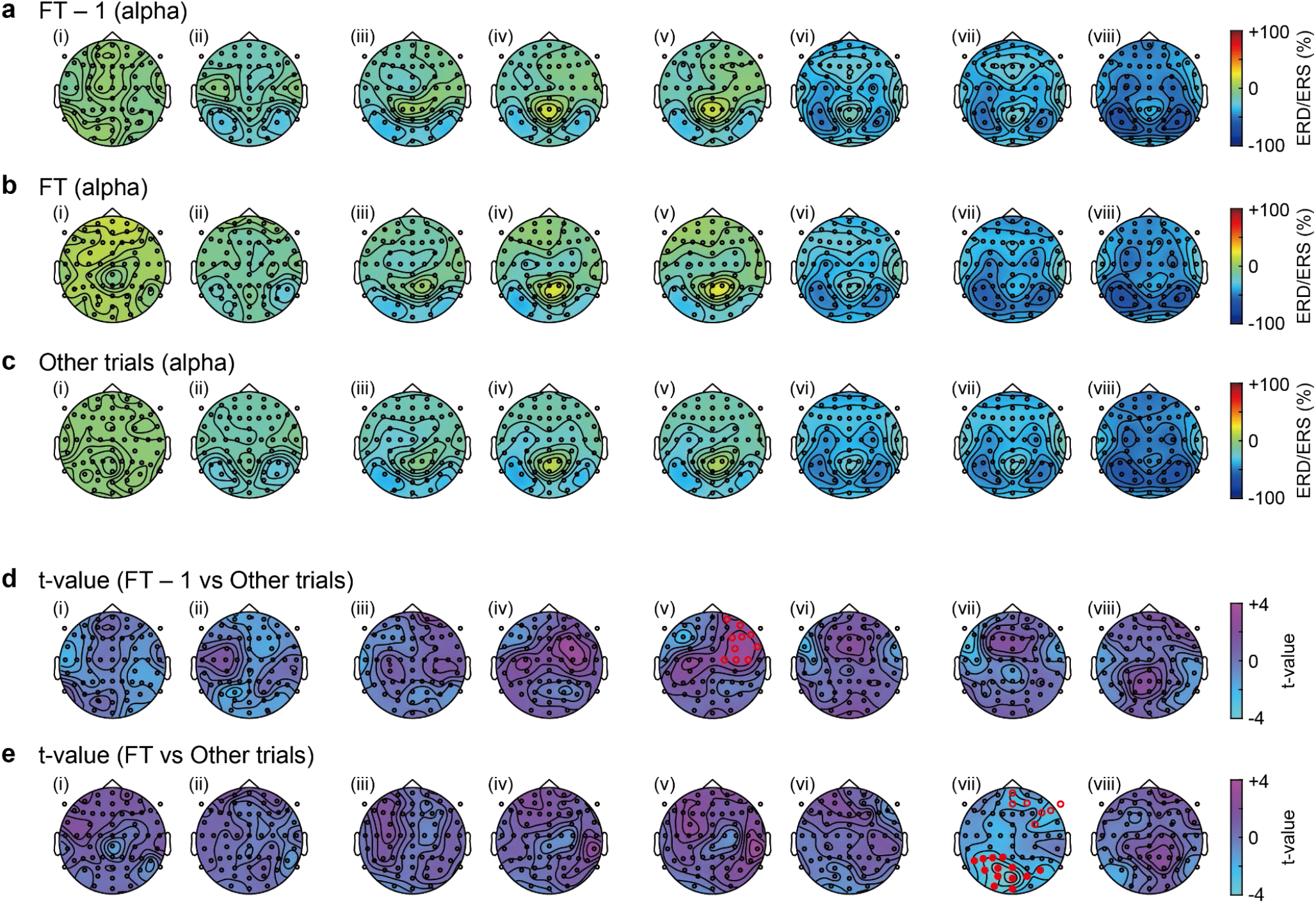
Topographic maps of the alpha ERD/ERS. (**a**) ERD/ERS in the trials preceding the FTs, (**b**) in the FTs, and (**c**) in the other trials averaged across subjects. i–viii indicate the time-epoch number in Fig. 3a. ERD/ERS in the FTs averaged across subjects. (**d**) In comparison with the ERD/ERS of the other trials, *t*-values obtained with paired *t*-tests with the trials preceding the FTs and (**e**) in the FTs are shown. Red rings denote *P* < 0.10 from cluster-based permutation testing. Red filled circles denote *P* < 0.05 from cluster-based permutation testing.

